# Defining the age-dependent and tissue-specific circadian transcriptome in male mice

**DOI:** 10.1101/2022.04.27.489594

**Authors:** Christopher A. Wolff, Miguel A. Gutierrez-Monreal, Lingsong Meng, Xiping Zhang, Lauren G. Douma, Hannah M. Costello, Collin M. Douglas, Elnaz Ebrahimi, Bryan R. Alava, Andrew R. Morris, Mehari M. Endale, G. Ryan Crislip, Kit-yan Cheng, Elizabeth A. Schroder, Brian P. Delisle, Andrew J. Bryant, Michelle L. Gumz, Zhiguang Huo, Andrew C. Liu, Karyn A. Esser

**Author notes:** These authors contributed equally (Christopher A. Wolff, Miguel A. Gutierrez-Monreal). Senior Author (Andrew J. Bryant, Michelle L. Gumz, Zhiguang Huo).

## Abstract

Cellular circadian clocks direct a daily transcriptional program that supports homeostasis and resilience. Emerging evidence supports age-associated changes in circadian functions. To define age-dependent changes at the systems level, we profiled the circadian transcriptome in the hypothalamus, lung, heart, kidney, skeletal muscle, and adrenal gland in 3 age groups. We found age-dependent and tissue-specific clock output changes. Aging reduced the number of rhythmically expressed genes (REGs), indicative of weakened circadian control. Many genes gained rhythmicity in old tissues, reflecting an adaptive response. REGs were enriched for the hallmarks of aging, adding a new dimension to our understanding of aging. Differential gene expression analysis found that there were temporally distinct clusters of genes in tissue-specific manner. Increased daily gene expression variability is a common feature of aged tissues. This novel analysis extends the landscape of the understanding of aging and highlights the impact of aging on circadian clock function and temporal changes in gene expression.

**HIGHLIGHTS:**

- Rhythmically expressed genes (REGs) in Young, but not Old mice, are enriched for the aging hallmarks across all tissues.
- The numbers of REGs decline across all tissues with age implicating the circadian clock in altered homeostasis.
- Age- and tissue-specific differentially expressed genes (DEGs) cluster at specific times of the day.
- Increase in gene expression variability over a day is a common feature of aging tissues.

## INTRODUCTION

Aging is characterized by a progressive loss of homeostatic control, leading to functional declines and decreased resilience. Over the last three decades, there have been numerous studies that used microarray and RNA-seq to examine age-related changes in gene expression across tissues (reviewed in (Frenk and Houseley, 2018). These studies contributed to the mechanistic understanding of aging biology, leading to defined hallmarks of aging (Kennedy et al., 2014; López-Otín et al., 2013). More recent comprehensive RNA-seq studies in rodents captured age-dependent transcriptomic changes across multiple organs and various ages and highlighted age-related increases in inflammation and loss of proteostasis across tissues (Schaum et al., 2020; Shavlakadze et al., 2019). However, these previous studies did not consider the time of day or the impact of aging on the circadian clock, thus overlooking a critical dimension of aging physiology.

In mammals, virtually every cell in the body has a functional circadian clock. The circadian system consists of a network of central and peripheral oscillators that give rise to various rhythmic outputs largely in a tissue-specific manner (Takahashi, 2017; Zhang et al., 2014). The suprachiasmatic nuclei (SCN) of the hypothalamus serve as the central clock that receives the daily light input and regulates the sleep/wake cycle (Mohawk et al., 2012). SCN neurons and peripheral tissue cells share a similar molecular clock mechanism which is based on an autoregulatory transcriptional negative feedback loop. The core feedback loop consists of the transcriptional activators BMAL1 and CLOCK and their negative regulators PER and CRY (Takahashi, 2017). The core clock also directs a daily transcriptional program that is cell typespecific. It is this circadian transcriptional output that prepares the cell for daily environmental changes and underlies predictive vs. reactive homeostasis (Koronowski and Sassone-Corsi, 2021; Moore-Ede, 1986). Recent studies have established that circadian functions decline over the lifespan. For example, age-related changes in the timing and amplitude of sleep/wake activity, body temperature, and hormone release in rodents and humans have been well documented (reviewed in (Hood and Amir, 2017)). Aging is also associated with a reduced ability to re-entrain to a new light/dark cycle, and increased mortality following repeated “jet lag” (Davidson et al., 2006; Inokawa et al., 2020; Sellix et al., 2012). It can be posited that circadian attenuation likely contributes to increased damage accumulation (Gladyshev et al., 2021), frailty phenotypes (Fried et al., 2021) and decreased resilience seen with aging (Kirkland et al., 2016).

Although age-related decline of circadian physiology and behavior has been generally recognized, significantly less is known about age-related changes in the circadian transcriptional output. A few recent aging-related circadian transcriptomic studies show that the core clock genes in aged tissues remain largely intact under light/dark conditions but the genes comprising the transcriptional output are altered (reviewed in (Welz and Benitah, 2020)). For example, the circadian transcriptome is reprogrammed from 3 months to 24 months in the mouse liver (Sato et al., 2017), and this was also apparent in stem cells from skin and muscle of 18-month-old mice (Solanas et al., 2017). Age-dependent circadian transcriptomic reprogramming has also been reported for the human prefrontal cortex (Chen et al., 2016) as well as in Drosophila (Kuintzle et al., 2017), demonstrating that altered circadian output is a conserved characteristic of aging. While these initial investigations brought attention to aging circadian clocks, there has been a lack of systematic design and analyses of the circadian transcriptome across organs and ages. We therefore carried out a 48 hour circadian transcriptomic analysis (Hughes et al., 2017) in male mice at 3 ages, (6, 18 and 27 months) and 6 organs and tissues (hypothalamus, lung, heart, kidney, skeletal muscle, and adrenal gland). We define age-related changes in the number and identity of the clock output. The temporal resolution of our data also offered an opportunity to examine the genes that were not rhythmic but displayed differential expression patterns at 4 distinct time domains of the day, (e.g., active phase vs. rest phase). We suggest that altered circadian clock output with age should be considered a hallmark of aging that contributes to the changes in cell and tissue homeostasis and likely contributes to frailty and compromised resilience in the old. To disseminate this data, we constructed a “CircaAge” database that provides investigators the ability to query the expression patterns of any gene in any of the organs across circadian time and age. Mechanistic insights into the interplay between the circadian and aging systems will offer new opportunities to enhance circadian function and promote healthy aging.

## RESULTS

### Profiling the aging circadian transcriptome across organs

Our study examined the circadian transcriptome in multiple tissues at multiple time points across the lifespan. We obtained male C57B6/J-NIA mice at 4, 16, and 25 months old (mo). The mice were maintained under 12h:12h light/dark conditions until 6 mo (Young), 18 mo (Aged), or 27 mo (Old) of age. Prior to tissue harvest, mice were released into constant darkness (circadian time or CT0) to study circadian gene expression under free-running conditions. Tissue collections began at CT18 and continued every 4 hours for 48h, with a total of 12 time points, in accordance with the guidelines for analysis of circadian genome data (Hughes et al., 2017). Our systems-level analysis included the hypothalamus, which contains the central SCN clock, and 5 other peripheral tissues, lung, kidney, skeletal muscle, heart, and adrenal gland (Figure 1A). We obtained high-quality RNAseq data from all tissues/organs with the exception of the first 24h of data from the adrenal gland of aged mice due to a technical issue. This data was included as supplemental data (Figure S1) but excluded in our larger analysis.

**Figure 1.**
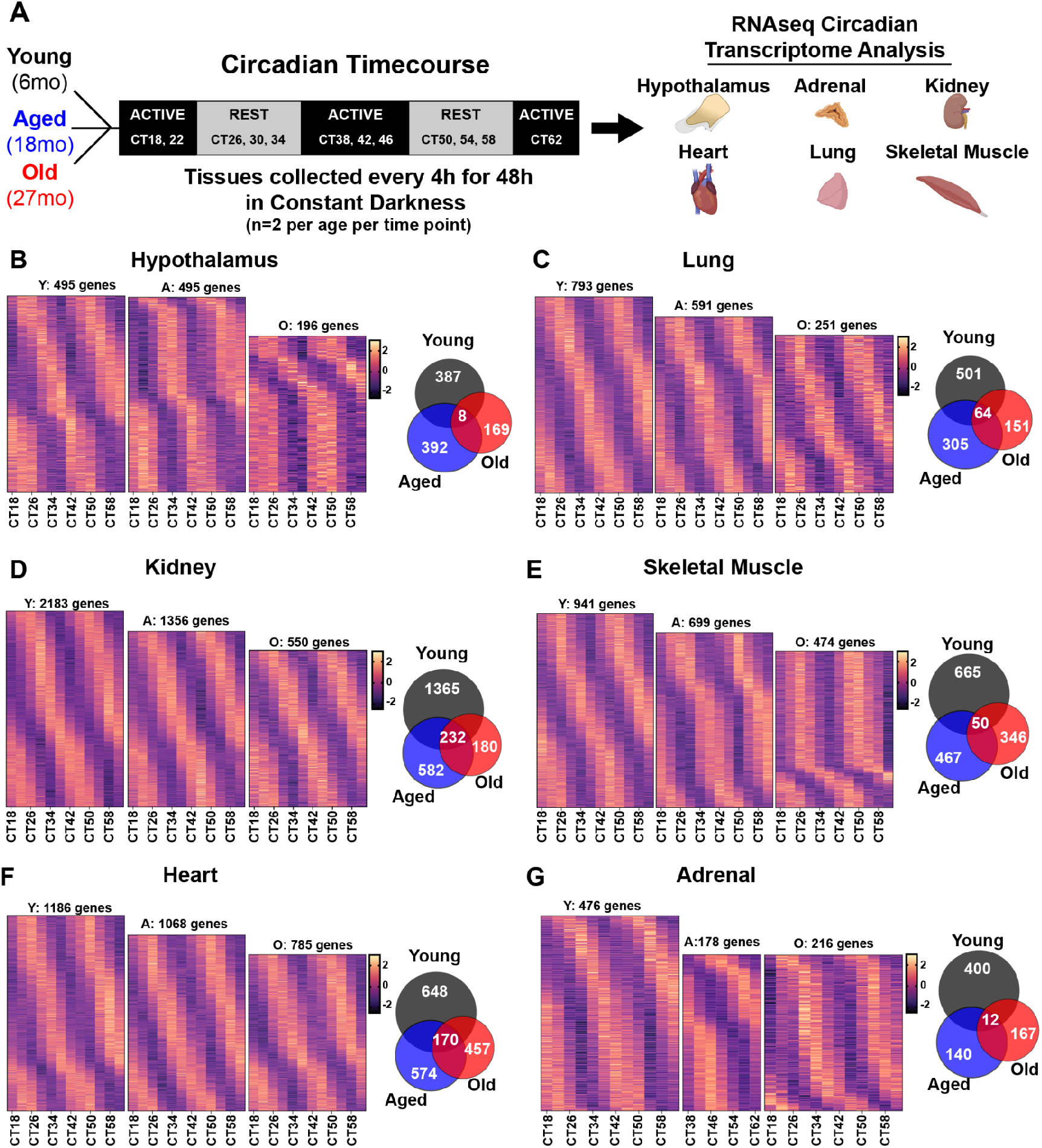
Circadian transcriptome analysis from 6 tissues and 3 ages. (A) Simplified study design schematic. Heat map of z-scored rhythmically expressed genes (REGs) and a Venn diagram of the REGs from each age for the (B) hypothalamus, (C) lung, (D) kidney, (E) skeletal muscle, (F) heart, and (G) adrenal gland. Due to technical difficulties with adrenal samples, only CT38-CT62 from the Aged were included for circadian transcriptome analysis.

### Age-dependent decline in the number of circadian genes across tissues

All RNAseq samples were sequenced to a depth of at least 40 million reads aligned to the mouse genome. To identify the circadian transcriptome, we deployed the cosinor model implemented in the diffCircadian software (Ding et al., 2021). Specifically, we defined circadian genes as those with 24h cosine oscillations in transcript abundance based upon a raw p-value < 0.01. We first examined the impact of age on the circadian rhythmically expressed genes (REGs) in each organ and tissue (Table S1). Overall, the majority of the REGs in the Young were tissue-specific, consistent with previous multiorgan genomic studies using young mice (Zhang et al., 2014). Across all tissues, we found an age-associated decline in the number of REGs. The largest REGs change was seen in the kidney with ~75% decline from Young to Old. The heart showed the least change with ~34% decline. The REGs within each tissue were also age-dependent, with less than 10% of REGs conserved across ages (Figure 1B-G and Figure S2). In the hypothalamus, there were 495 REGs in both the Young and Aged groups, but this declined to 196 in Old (Figure 1B and Table S2). We identified 793 REGs in the Young lung, 591 in Aged, and 251 in Old (Figure 1C and Table S3). The kidney had 2,183 REGs in Young, 1,356 in Aged, and 550 in Old (Figure 1D and Table S4). As in the hypothalamus, the kidney circadian transcriptome was clearly age-specific, with only 232 (8%) of the REGs shared across all three ages. In skeletal muscle, the number of REGs decreased from 941 in Young to 699 in Aged and 474 in Old (Figure 1E and Table S5). Again, there was limited overlap of the circadian transcriptome across ages, with just 50 (3%) of the REGs shared across all three ages. We identified 1,186 REGs in Young hearts, 1,068 in Aged, and 785 in Old, with 170 genes shared across all three ages (Figure 1F and Table S6). Finally, in the adrenal gland, we identified 476 REGs in Young, and only 178 in Aged and 216 in Old (Figure 1G). Our analyses demonstrated that the number of REGs declines in all tissues in an age-dependent manner, and that few genes are rhythmically expressed across age.

With such large changes in the circadian output with age, we queried the aging effects on the expression patterns of the core clock genes representing the three interlocking loops of the circadian clock (Figure S3). The expression of *Bmal1* was most robust with age being largely rhythmic across all tissues and ages. The one exception was that the core clock genes in the hypothalamus were either weakly rhythmic or not rhythmic across all ages, likely due to the differences in circadian timing across the cell types and nuclei in this brain region (Wen et al., 2020; Zhang et al., 2014). Recent studies had suggested a limited impact of age on the core clock genes (Sato et al., 2017; Solanas et al., 2017). However, our data revealed a notable decline in the rhythmicity of the repressor components of the clock. We found that *Per1, Per2, Cry1*, and *Cry2* were dampened from Aged to Old across all peripheral tissues (Figure S3). The secondary loop genes, *Nr1d1* and *Nr1d2*, also exhibit age-related loss of rhythmicity but only in the skeletal muscle and hypothalamus. Thus, there are common aging effects on the negative limb components of the core clock mechanism across tissues. This observation is consistent with the slowed rate of entrainment of the circadian system with age (Sellix et al., 2012).

### Temporal patterns of age-associated differential gene expression

Our time course collection also provided the opportunity to explore temporal differences in age-related changes in gene expression beyond circadian rhythmicity. We binned the 48h data over 4 time domains or phases: the rest phase (light phase in the nocturnal mice), activity onset (rest-active transition), active phase (dark phase), and activity offset (active-rest transition). We used an ordinal analysis strategy to define time domain specific genes that changed in the same direction from Young to Aged to Old. This approach is similar to the linear gene expression changes noted by Shavlakadze and colleagues (Shavlakadze et al., 2019). To our surprise, we found that a large number of the differentially expressed genes (DEGs) were detected at unique time domains but this occurred in a tissue-specific manner. These outcomes provide unique time of day molecular maps of tissue aging with the potential to more precisely target therapeutic strategies. We describe in each of the sections below the unique clusters of DEGs for the distinct time domain.

### Loss of circadian metabolic and immune homeostasis in the aging hypothalamus

The hypothalamus controls essential homeostatic and survival-related functions. It is a small yet highly heterogeneous tissue, with multiple cell types and functionally distinct nuclei, including the SCN. Through projections to other nuclei, the central SCN clock functions to coordinate various circadian rhythms such as endocrine release and sleep/wake behavior (Acosta-Rodríguez et al., 2021; Kramer et al., 2022). The RNA-seq sensitivity enabled the detection of cellular markers specific to cell types and nuclei, for example, *Vip* and *Nms* (SCN), *Agrp, Npy* and *Pomc* (arcuate nucleus), *Hcrt* (lateral hypothalamic area), *Gfap* (astrocytes), and *Aif1* (microglia) (Chen et al., 2017). The high-quality data provide the first systems-level view of circadian transcripts across ages in the hypothalamus. Circadian rhythmicity analysis uncovered 495 REGs in Young and Aged, but significantly less (196) in Old. Overall, there were fewer REGs in the hypothalamus than in the peripheral tissues at the same cutoff, which is due in part to its cellular heterogeneity (Wen et al., 2020). While some REGs were present in all three ages, most of them were specific to one age group. Most REGs in Young lost rhythmicity in Old, whereas several genes gained rhythmicity in Old.

We used the Ingenuity Pathway Analysis (IPA) to analyze and integrate the functional pathways enriched for the REGs across ages. This analysis revealed conserved pathways across ages as well as age-specific pathways (Figure 2A-B and Table S7). For example, the NRF2 oxidative response is rhythmic across the lifespan, although the genes contributing to the pathway were different across ages. The HIF1, adipogenesis, and neuregulin signaling pathways lost REG enrichment from Young and Aged to Old (Figure 2A). Strikingly, many pathways were enriched only in one of the three ages, with the highest number of the agespecific pathways occurring in Young (n=80), followed by Aged (n=37) and Old (n=27) (Figure 2B). Interestingly, many of these REG pathways in Old emerge from unique gain-of-rhythm genes. In general, the REG pathways overrepresented in Young were involved in circadian regulation, energy homeostasis, proteostasis, and cell growth and development, whereas those in Old were related to stress and adaptive responses and basic central nervous system functions such as neurogenesis, axonogenesis, and synapse formation.

**Figure 2.**
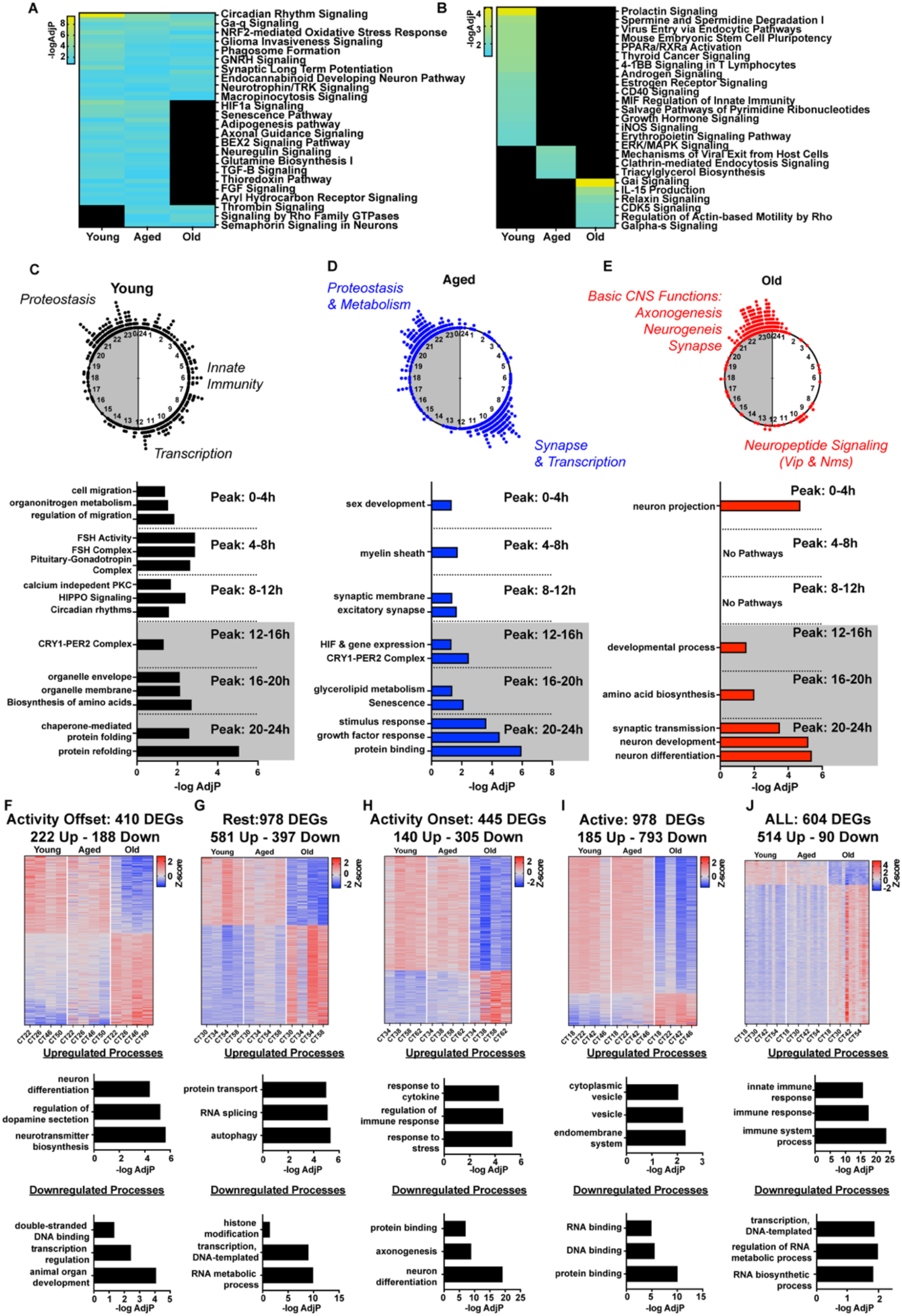
Age-dependent circadian transcriptomic changes in the hypothalamus. (A) Shared IPA pathways enriched by the REGs that cycle in 2-3 age groups. (B) Age-specific circadian oscillatory IPA pathways. (C-E) Peak time maps of all oscillating genes from the hypothalamus of (C) Young, (D) Aged and (E) Old mice. Each dot represents the peak time of a single significantly circadian gene. Underneath the peak time map is a histogram of top time of day specific gene expression pathways. (F-I) Heat map of z-scored age-related differentially expressed genes (DEGs) and histograms of up- or down-regulated pathways from the specific gene sets from the active to rest transition period (F), the rest phase (G), the rest to active transition period (H), and the active period (I). (J) Heat map of z-scored age-related DEGs from all time domains and histograms of up- or down-regulated pathways from the specific gene sets.

We next analyzed the REGs based on their circadian time (CT) of peak expression for each age group (Figure 2C-E, top panels). Strikingly, we observed an age-specific daily distribution of the REGs. While the REGs in the Young hypothalamus had multiple peaks across the circadian cycle, only two peaks were prominent in Aged, with one activity offset and the other at activity onset. The top enriched functional pathways reflect the time of day-dependent hypothalamic functions (Figure 2C-E, bottom panels and Table S8). Notably, the REGs in the CRY1-PER2 complex had a Peak Time between 12-16h in Young and Aged lost their rhythms in Old, indicative of compromised active circadian repression and weakened transcriptional outputs. Among the genes that were not highly rhythmic in Young or Aged but gained robust rhythms in Old were vasoactive intestinal polypeptide (*Vip*) and neuromedin S (*Nms*). These two neuropeptides play key roles in photic entrainment, neuronal coupling, and SCN synchronization. Other examples include the corticotropin-releasing hormone (*Crh*) in the hypothalamus-pituitary-adrenocortical (HPA) axis and factors involved in neuropeptide signaling (e.g., *Auts2, Drd2, Chrm3*, and *Adora2a*), which became highly rhythmic in Old, likely as an adaptive response to homeostatic stress.

In addition to the REGs, we found a large number of DEGs in the hypothalamus that exhibited unique expression patterns across the 4 time domains of the day (Figure 2F-J and Table S9). There were 604 DEGs that were either up-regulated (514) or down-regulated (90) across ages at all time domains (Figure 2J). Pathway analysis revealed that the upregulated DEGs had an overrepresentation for the immune and inflammatory responses in Aged and Old tissues (e.g., *Gfap, Aif1, Trem2, Adgre1*, and *Ptgs1*), whereas active transcription-related genes were significantly downregulated. The shift from the metabolically active and proliferative state in Young to an inflammatory state in Old is consistent with findings from genomic studies in both mice (Hammond et al., 2019; Schaum et al., 2020) and in rats (Shavlakadze et al., 2019). The DEGs at activity offset were enriched in neurotransmitter synthesis and neuron differentiation and those at activity onset were in stress and immune responses. Also of note, during the rest phase, genes related to protein transport (e.g., *Vps35, Vps39, Vps45*) and autophagy (e.g., *Atg3*) were significantly upregulated with age (Figure 2G), whereas those involved in vesicular transport (e.g., *Srebf1, Slc2a8, Tbc1d17*) were upregulated during the active phase (Figure 2I). Taken together, our findings from both the REGs and the DEGs highlight the importance of considering the time of day when exploring age-related changes in gene expression and functions.

### Inflammation is a major feature of aging in the lung

Broadly speaking the lung functions primarily for gas exchange, however, the organ also serves as a first-line site for defense against pathogenic microbial species (Eddens and Kolls, 2012; Skloot, 2017). Therefore, time of day responses to stressors are critical for maintaining healthy pulmonary and organismal function. The lung REGs were enriched for several stress-related pathways that were maintained across the lifespan, including the unfolded protein response and xenobiotic metabolism (Figure 3A and Table S10). However, immune-related pathways, such as CXCR4 signaling and the hypoxia pathway, HIF1a, lost rhythmicity in the Old (Figure 3B). For example, *Tmem173/Sting1* and *Unc93b1* were robustly rhythmic in the Young but lost rhythmicity with age. The REG pathways that gained rhythmicity in Old were enriched for amino acid metabolism and amino acid hormone synthesis (Figure 3B).

**Figure 3.**
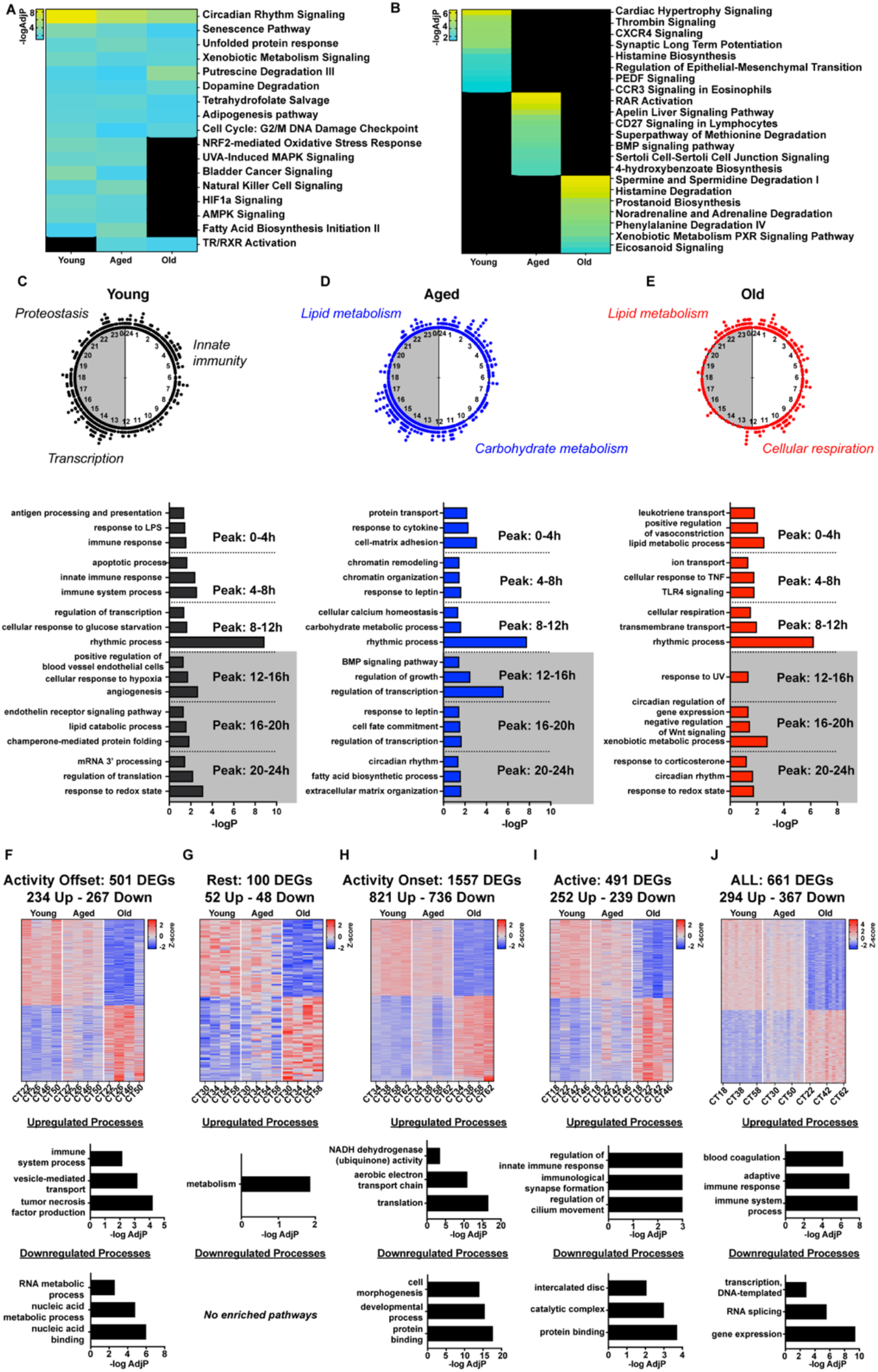
Age-associated changes in the lung circadian transcriptional output and time of day dependent interpretations of age-related changes in gene expression. (A) Shared IPA pathways enriched by the REGs that cycle in 2-3 age groups. (B) Age-specific circadian oscillatory IPA pathways. (C-E) Peak time maps of all oscillating genes from the lungs of Young (C), Aged (D) and Old mice (E). Each dot represents the peak time of a single significantly circadian gene. Underneath the peak time map is a histogram of top time of day specific gene expression pathways. (F-I) Heat map of z-scored age-related differentially expressed genes (DEGs) and histograms of up- or down-regulated pathways from the specific gene sets from the active to rest transition period (F), the rest phase (G), the rest to active transition period (H), and the active period (I). (J) Heat map of z-scored age-related DEGs from all time domains and histograms of up- or down-regulated pathways from the specific gene sets.

The peak time distribution of the REGs in the lung showed similarities across ages. However, there were significant age-specific changes in the temporal distribution of the REG functions (Figure 3C-E and Table S11). For example, in the early rest phase (Peak Time 0-8h), the REGs in the Young contribute to the immune response and antigen processing pathways but these pathways were lost in the Aged and Old lungs. This would implicate the diminished ability of the lungs to best defend against the immune challenges with age. In the Old, the pathways over-represented in the active part of the day (Peak Time 20-24h) include response to corticosterone and redox state suggesting a more reactive set of REG functions related to stress responses.

The lung exhibited large differences in the magnitude of the DEGs across the 4 time domains. Specifically, activity onset had the largest number of DEGs (1,557) while the rest phase showed the smallest number (100) (Figure 3F-I and Table S12). Oxidative metabolic pathways were up-regulated in the activity onset as well as the rest phase while immune-related DEGs were up-regulated in the active phase and activity offset. The up-regulation of immune response DEGs in the active phase suggests compensation for the loss of immune REGs in the Young. Overall, this reinforces a decrease in the anticipatory nature of the pulmonary response to pathogens, resulting in a time of day vulnerability to pulmonary disease. Thus, the aging-associated loss of circadian functions likely potentiates the inflammatory response in the lung, potentially contributing to loss of organismal resilience.

### Aging blunted the oscillatory expression patterns of ion transporters in the kidney

The kidney is critical for several aspects of systemic homeostasis (Seifter, 2019; Verschuren et al., 2020). One of its primary functions is to maintain electrolyte balance through ion transport. Relative to the other tissues, the kidney had the most REGs at Young (2,183) and Aged (1,356) but REG numbers decreased to 550 in Old. Functional cluster analysis of the REGs revealed that many pathways were maintained across age including the unfolded protein response and fibroblast growth factor (FGF) signaling (Figure 4A and Table S13). However, the aldosterone signaling pathway was enriched in the Young and Aged but lost in the Old (Figure 4A). Notably, the two key REGs unique to kidney function, *Scnn1a* (a subunit of the sodium channel) and *Atp1a1* (a subunit of the Na+/K+ -ATPases) were highly rhythmic in the Young but attenuated in the Old. The REG pathways were representative of the hallmarks of aging, including AMPK signaling and ubiquitination, lost oscillations in Old (Figure 4A). Of the pathways enriched solely in Aged, eNOS signaling likely reflects daily maintenance and homeostasis of the epithelium as well as renal hemodynamics (Nishimura et al., 2021) (Figure 4B). Finally, NAD^+^ salvage pathways and acute immune responses were unique to Old, possibly as an adaptive mechanism to maintain homeostasis.

**Figure 4.**
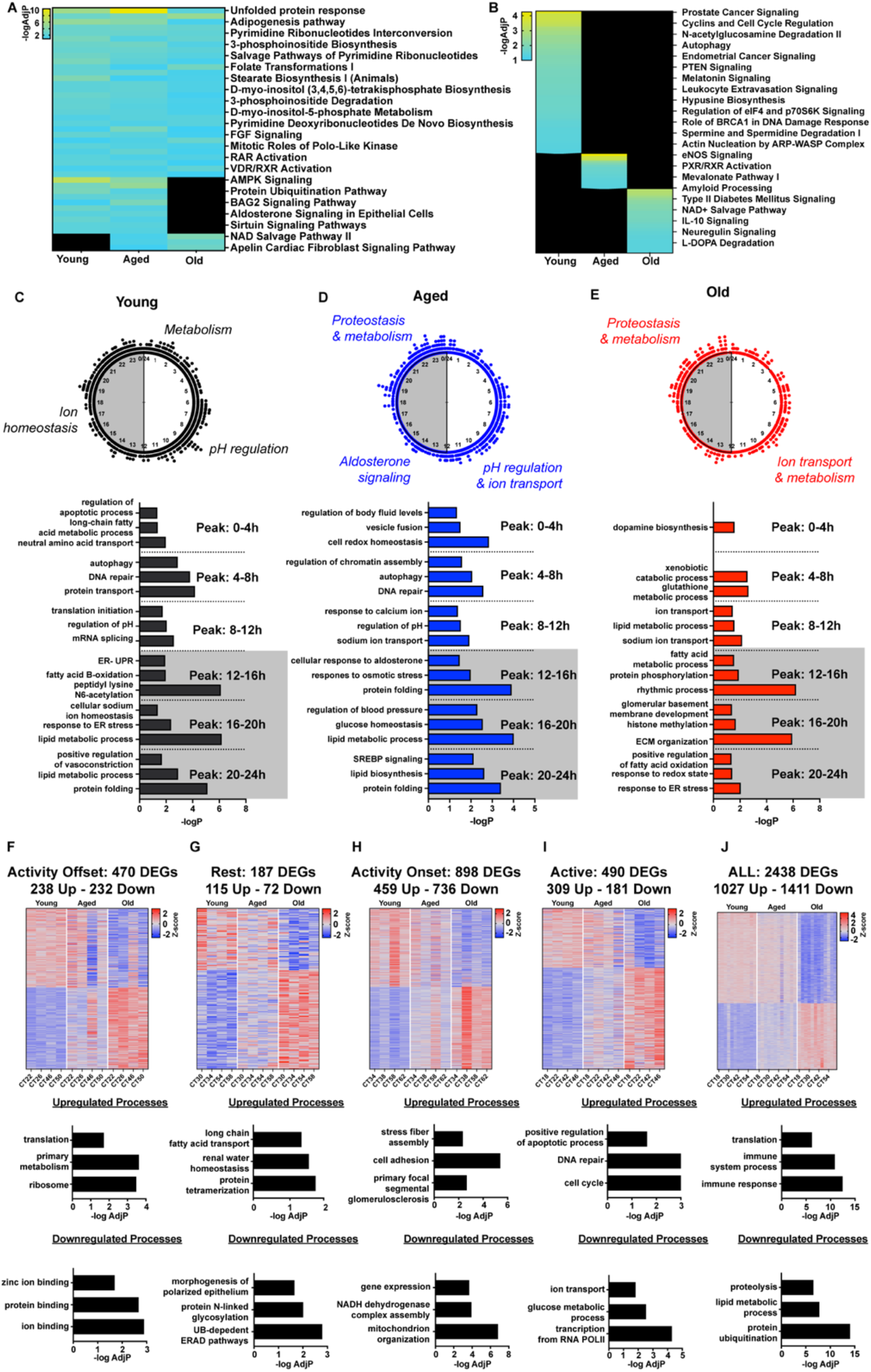
Age-associated changes in the kidney circadian transcriptional output and time of day dependent interpretations of age-related changes in gene expression. (A) Shared IPA pathways enriched by the REGs that cycle in 2-3 age groups. (B) Age-specific circadian oscillatory IPA pathways. (C-E) Peak time maps of all oscillating genes from the kidneys of Young (C), Aged (D) and Old mice (E). Each dot represents the peak time of a single significantly circadian gene. Underneath the peak time map is a histogram of top time of day specific gene expression pathways. (F-I) Heat map of z-scored age-related differentially expressed genes (DEGs) and histograms of up- or down-regulated pathways from the specific gene sets from the active to rest transition period (F), the rest phase (G), the rest to active transition period (H), and the active period (I). (J) Heat map of z-scored age-related DEGs from all time domains and histograms of up- or down-regulated pathways from the specific gene sets.

The overall peak time distribution of the REGs in the kidney did not change dramatically with age. However, pathway analysis highlighted age-specific changes in the temporal distribution of the REG functions (Figures 4C-E and Table S14). For instance, in the Young and Aged kidneys, genes related to pH homeostasis were oscillating with a Peak Time between 8-12h, but not in Old. While ion transport processes remained rhythmic in the Old, they peaked at different times compared to Young and Aged. Another notable example was proteostasis-related pathways that were enriched across the circadian cycle in Young and Aged but limited to the late active phase in Old. While pH homeostasis and proteostasis related genes were rhythmic in the Young and Aged kidney, the daily timing becomes out of phase with age. These age-dependent changes in temporal alignment between key physiological processes may exacerbate aging kidney phenotypes.

There were profound age-related changes in the DEGs, with 2,438 genes differentially regulated with age at all times, in addition to the DEGs that were specific to different time domains (Figure 4F-J and Table S15). We observed downregulation of mitochondrial genes and an increase in stress fiber assembly at activity onset. Moreover, the upregulated genes at activity onset were enriched for primary focal segmental glomerulosclerosis, highlighting kidney-specific changes in gene expression with age (Figure 4H). As in the lung and hypothalamus, there was a significant upregulation of immune-related pathways with age at all time domains, accompanied by a decrease in proteolysis and lipid metabolism (Figure 4J). Additionally, the decreased proteostasis REGs in Old were paralleled by the decreases in the DEGs (Figure 4F-J; bottom panels).

### Skeletal muscle circadian transcriptome changes with age highlight autophagy and myogenic programs

Skeletal muscle is critical for health through its role in regulating movement and metabolism and it is emerging as a source of circulating factors such as myokines (Cartee et al., 2016; Wolfe, 2006). In the context of aging, epidemiological studies have shown strong correlations between loss of muscle strength and increased morbidity and mortality (REF?). As with other tissues, there was a significant decline in the number of REGs with age. Across the REGs there was a preservation of functional groups including insulin receptor and HIF1 signaling, but pathways such as AMPK, PI3K/AKT and unfolded protein response lost rhythmicity with age (Figure 5A and Table S16). The functional categories that gained rhythmicity with age include dilated cardiomyopathy and IL-1 signaling (Figure 5B).

**Figure 5.**
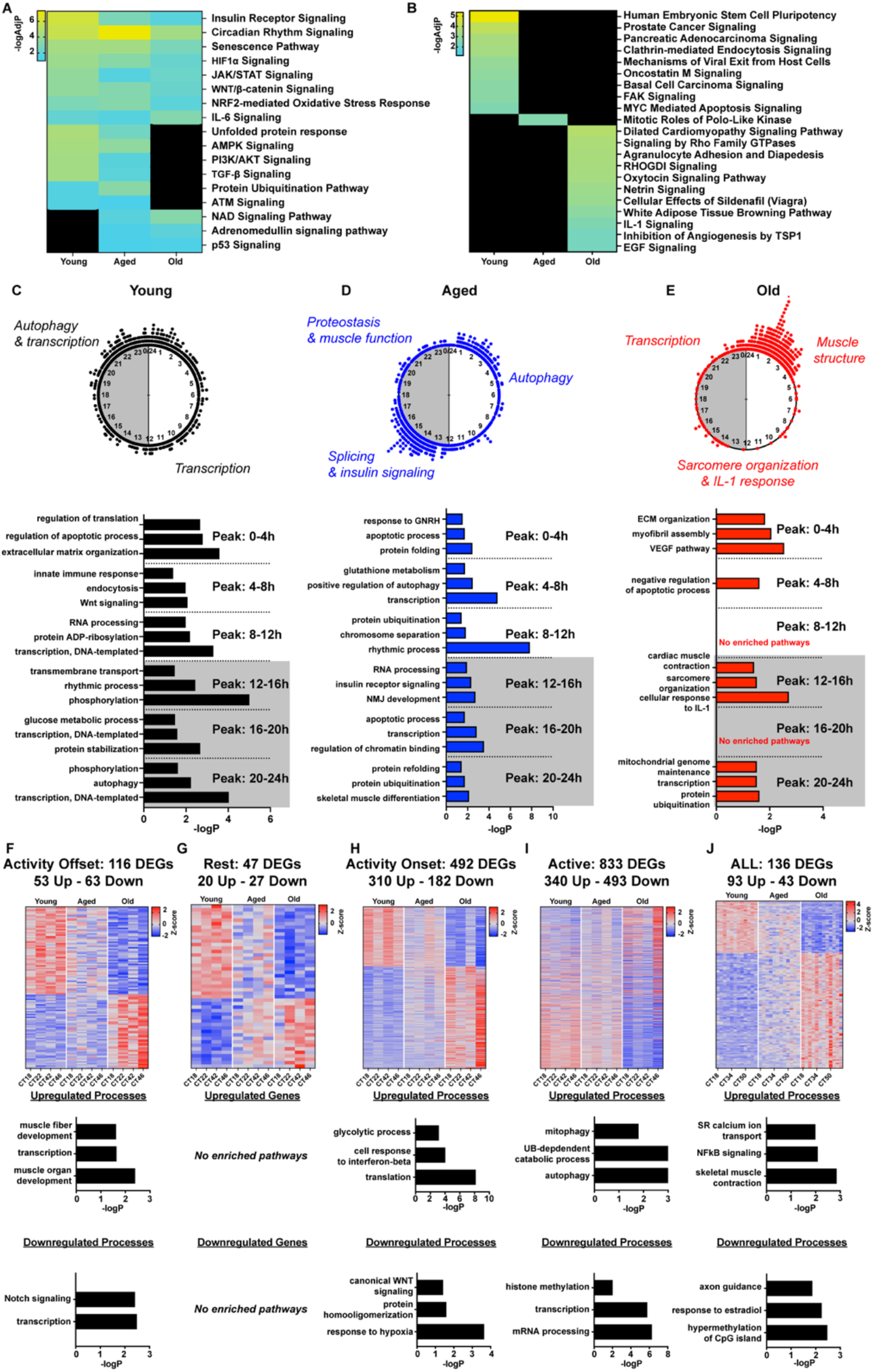
Age-associated changes in the muscle circadian transcriptional output and time of day dependent interpretations of age-related changes in gene expression. A) Shared IPA pathways enriched by the REGs that cycle in 2-3 age groups. (B) Age-specific circadian oscillatory IPA pathways. (C-E) Peak time maps of all oscillating genes from the skeletal muscle of Young (C), Aged (D) and Old mice (E). Each dot represents the peak time of a single significantly circadian gene. Underneath the peak time map is a histogram of top time of day specific gene expression pathways. (F-I) Heat map of z-scored age-related differentially expressed genes (DEGs) and histograms of up- or down-regulated pathways from the specific gene sets from the active to rest transition period (F), the rest phase (G), the rest to active transition period (H), and the active period (I). (J) Heat map of z-scored age-related DEGs from all time domains and histograms of up- or down-regulated pathways from the specific gene sets.

The distribution of the peak times for the REGs in muscle was similar to that in the hypothalamus. In the Young the REGs were distributed across the day, but in the Aged the REGs were largely bimodal with clusters at activity onset and offset (Figure 5C-E and Table S17). Of note, the REGs that contribute to autophagy peaked at activity offset in Young and *Tfeb*, considered a key upstream regulator of autophagy (Napolitano and Ballabio, 2016; Settembre et al., 2011), is one of the REGs in that temporal cluster. In the Aged, *Tfeb* is no longer circadian and the autophagy cluster is centered toward the middle of the rest phase (Figure 5D). Recent studies have implicated changes in the timing of autophagy as a contributor to aging in flies and mice (Juste et al., 2021; Ulgherait et al., 2021). The REGs in the Old muscle were largely unimodal, peaking at activity offset. This cluster includes myofibril assembly and VEGF pathway suggesting enrichment of muscle tissue maintenance functions in the Old (Figure 5E).

Analysis of the muscle DEGs revealed significant differences across the 4 different time domains. The largest cluster of DEGs was found in the active phase (833 DEGs) with only 47 DEGs in the rest phase and 136 DEGs were found in all time domains (Figure 5F-J and Table S18). In the active phase, the down-regulated DEGs contributing to RNA processing, transcription, and genome maintenance were overrepresented, whereas autophagy and mitophagy pathways were up-regulated DEGs with age. The increase in autophagy DEGs may compensate for the loss of rhythmic control of autophagy with age. Among the DEG pathways that were common across all time domains was the up-regulated NF-kB signaling, which is consistent with inflammation being a common issue that all tissues are responding to with age (Figure 5J). The other up-regulated cluster across all time domains was related to skeletal muscle contraction and sarcoplasmic reticulum (SR) calcium transport. The age-related change in the myogenic program is consistent with an increased, albeit noisy, expression of the myogenic regulatory factor *Myf6* seen with age.

### Limited impact of age on circadian and steady-state mRNA expression in mouse hearts

The heart is required for maintenance of gas and nutrient delivery as well as cellular waste removal through the circulation (Pittman, 2013; Rana et al., 2020). The heart is unique among all the tissues in that the core clock components exhibit very little change with age and the REG pathways were largely conserved across age. These include many pathways linked to the hallmarks of aging such as protein homeostasis and senescence (Figure 6A and Table S19). There were some pathways enriched only in the Young and Aged, including some essential processes like p53 and iNOS signaling that were no longer enriched in the Old. In contrast to the other tissues, the heart does not have new REG pathways unique to the Old. (Figure 6B).

**Figure 6.**
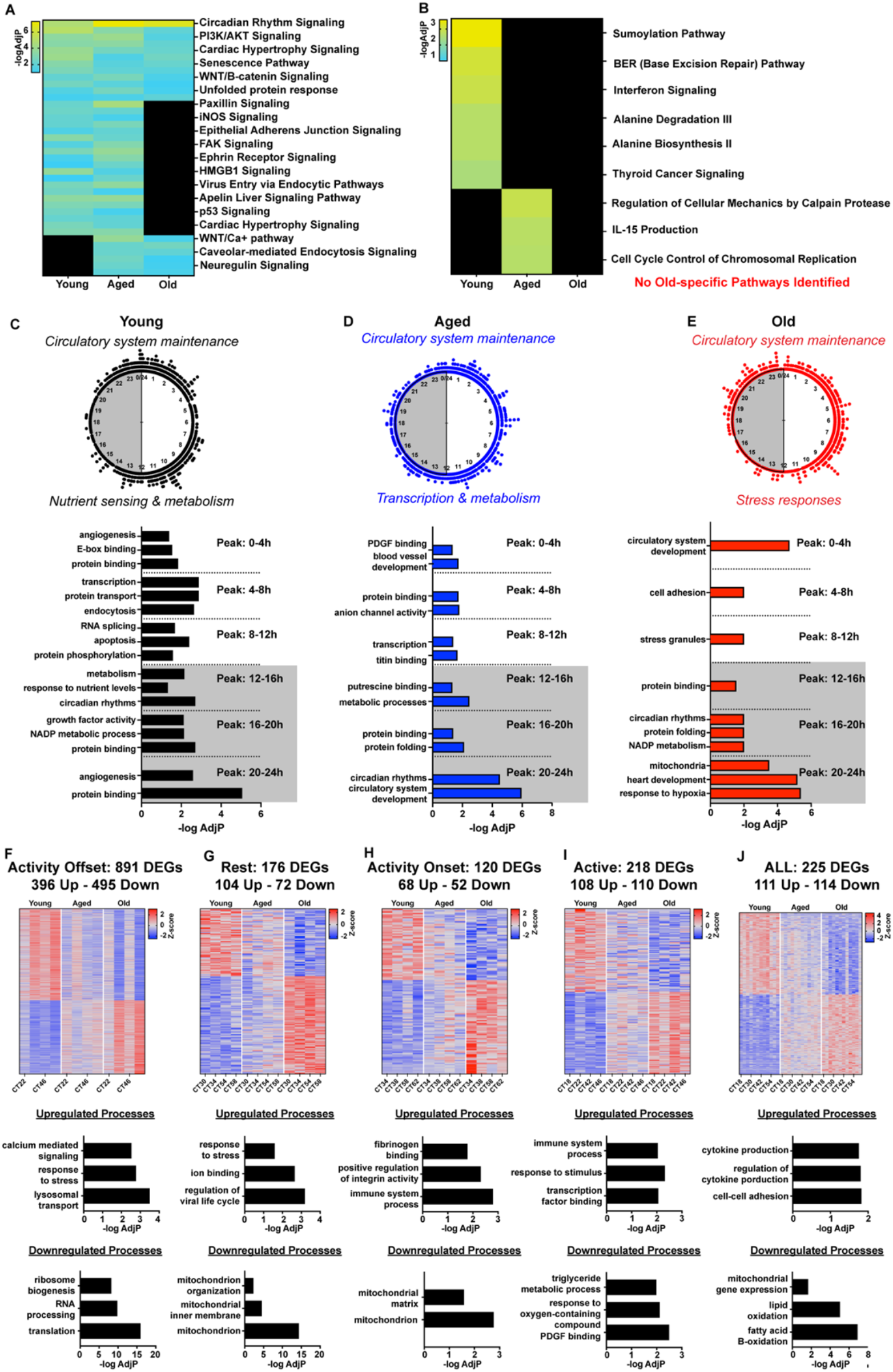
Age-associated changes in the heart circadian transcriptional output and time of day dependent interpretations of age-related changes in gene expression. A) Shared IPA pathways enriched by the REGs that cycle in 2-3 age groups. (B) Age-specific circadian oscillatory IPA pathways. (C-E) Peak time maps of all oscillating genes from the heart of Young (C), Aged (D) and Old mice (E). Each dot represents the peak time of a single significantly circadian gene. Underneath the peak time map is a histogram of top time of day specific gene expression pathways. (F-I) Heat map of z-scored age-related differentially expressed genes (DEGs) and histograms of up- or down-regulated pathways from the specific gene sets from the active to rest transition period (F), the rest phase (G), the rest to active transition period (H), and the active period (I). (J) Heat map of z-scored age-related DEGs from all time domains and histograms of up- or down-regulated pathways from the specific gene sets.

The peak time distribution of the REGs showed little change, consistent with very little apparent cardiac-specific circadian disruption. Across the three ages, the REGs were evenly distributed throughout the 24h day (Figure 6C-E and Table S20) similar to the kidney and lung tissues. In addition to maintaining this daily distribution of genes, pathways related to circulatory maintenance (e.g., angiogenesis and circulatory system development) were consistently peaking at the activity offset across all ages. Despite this pathway conservation with age, there were age-related changes in the heart REGs at the activity onset, where the Young and Aged hearts were enriched for metabolic processes, while the Old were enriched for stress responses.

While aging had a limited impact on the REGs in the heart, it had the highest number of unique DEGs at activity offset (891; Figure 6F). During this transition period, we observed age-related decreases in ribosome biogenesis, RNA processing, and translation processes, suggesting reduced proteostasis in the Old. Relevant to cardiac function, calcium signaling pathways were up-regulated at activity offset with age. In contrast, DEGs enriched for mitochondria and fatty acid oxidation were decreased at all time domains, supporting the concept of age-related decrements in metabolic capacity in the Aged and Old (Barton et al., 2016). Additionally, cytokine-related genes were upregulated with age across all time domains in the heart (Figure 6F-J and Table S21).

### Increased daily transcriptional variability is conserved across tissues

Most recently, studies have identified consistent age-associated increases in the variability of gene expression (Bahar et al., 2006; Enge et al., 2017). We queried the non-circadian genes and found that there was an increase in the variability of gene expression with aging in all tissues except the heart. For example, the hypothalamus and lung had 2,516 and 2,452 genes with high variation, respectively. Surprisingly, we found that even common housekeeping genes, including *Gapdh* and *Rplp0*, were significantly more variable with age across tissues (Table S22). Comparing the variable genes across tissues identified that 3 peripheral tissues, lung, kidney and skeletal muscle, shared 212 genes suggesting common age associated changes in transcriptional control (Figure 7G). Functional cluster analysis of those 212 variable genes identified protein transport and ubiquitination, chromatin organization, and immune system processes (Figure 7H and Table S22). The increased variability in gene expression is consistent with the concept that the physiology underlying frailty with age occurs through increased dysregulation (Fried et al., 2021).

**Figure 7.**
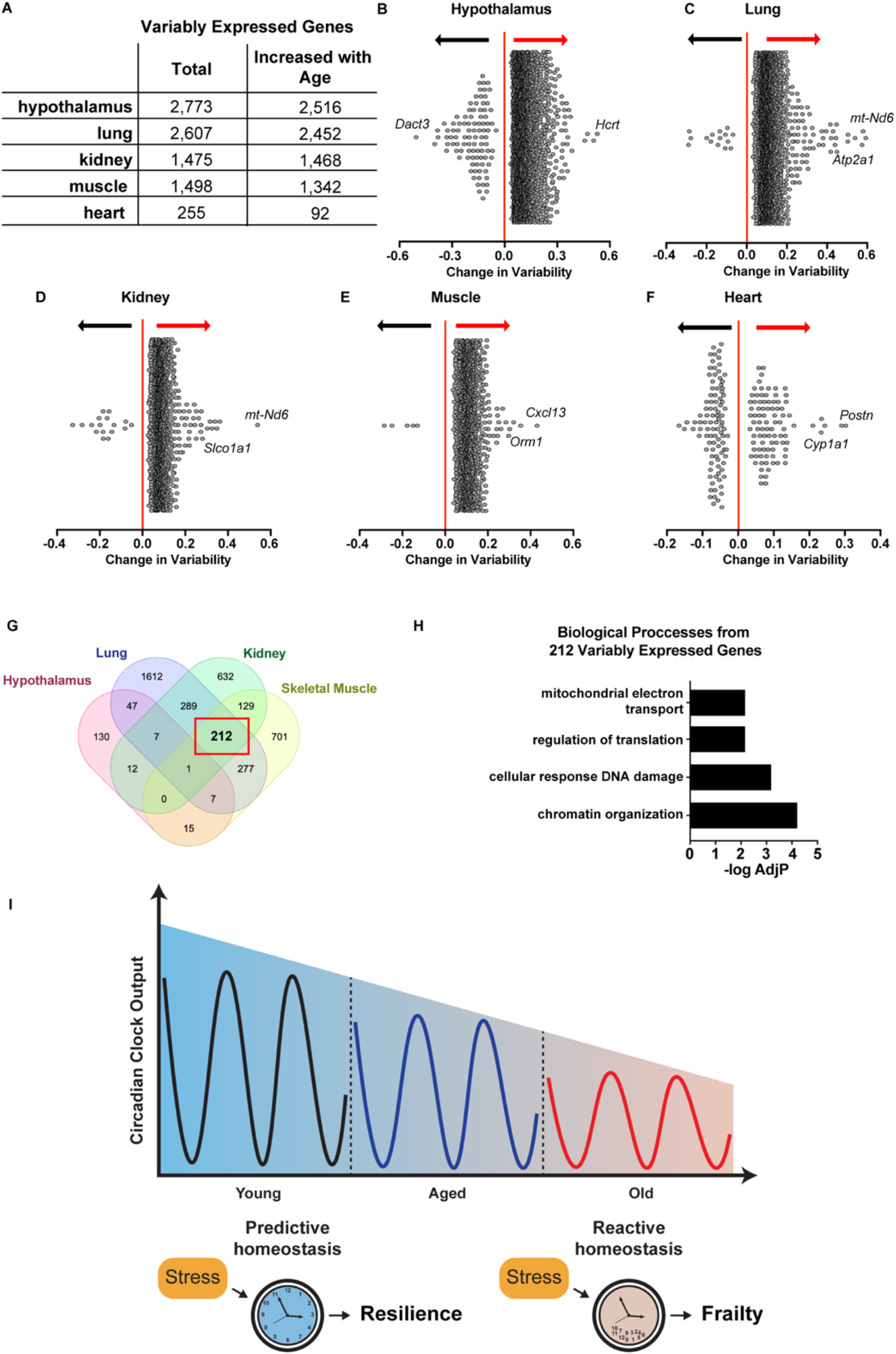
Age-related changes in variably expressed genes across tissues and summary of age-related changes in circadian gene expression. (A) Tissue-specific table of age-related changes in variable gene expression. (B) Hypothalamic variably expressed genes. “Change in Variability” means “change in variability (measured by absolute deviance) per unit change in age group (Young -> Aged or Aged -> Old)”. Change in Variability above 0 indicates genes that are more variably expressed with age, notated by the red arrow. (C) Lung variable gene dispersion plot (D) Kidney variable gene dispersion plot (E) Skeletal muscle variable gene dispersion plot (F) Heart variable dispersion plot (G) Venn diagram of overlapping variable genes across tissues (H) Top biological processes enriched by overlapping variable genes in the muscle, kidney, and lung and (I) Summary figure highlighting the age-related changes in circadian clock output (number of REGs) and a conceptual model demonstrating the age-related loss of predictive homeostatic control, likely contributing to frailty phenotypes.

### CircaAge: Database of Age-Dependent Changes in Circadian and Non-Circadian Gene Expression Patterns

Genome-wide age-dependent changes in gene expression, circadian or not, are difficult to visualize. To disseminate this data, we developed a web-application database, “CircaAge” (https://circaage.shinyapps.io/circaage/). This publicly accessible resource allows the user to query any gene(s) of their interest and visualize transcript expression patterns across 2 circadian days and 3 age groups in 1 central and 5 peripheral tissues. This database also provides model fitness parameters and significance levels of circadian rhythmicity. Users may specify (1) single or batch entry of genes of interest, (2) 1-3 ages of interest, and (3) 1-6 tissue types of interest. Data can be conveniently exported as *.csv files with statistical outputs including peak time, amplitude, basal expression (i.e., MESOR), phase, R^2^, and p-values (Ding et al., 2021).

## DISCUSSION

Aging is characterized by declines in physiological functions and a reduced capacity to maintain homeostasis (Kennedy et al., 2014; López-Otín et al., 2013; Pomatto and Davies, 2017). Aging is also characterized by declines in circadian functions, such as sleep/wake cycles, (Fonseca Costa and Ripperger, 2015; Hood and Amir, 2017; Zhao et al., 2019). The circadian clock within each cell directs a daily transcription program that temporally segregates important cell functions to support homeostasis and resilience. This clock function underlies what is known as predictive homeostasis as the changes within the cell and system occur prior to, and not in reaction to, known changes in the environment linked with the light/dark and rest/ activity cycles (Dibner and Schibler, 2015; Kim et al., 2019; Moore-Ede, 1986). Previous studies of circadian transcriptomes with age have been limited to the liver and stem cells and they were done in only two age groups (Sato et al., 2017; Solanas et al., 2017). Our goal was to obtain a systems-level understanding of the aging circadian transcriptome through analysis of 6 tissues and across 3 ages. In the present work, we identified significant changes in the circadian transcriptomes (i.e. REGs) with age and across all tissues. While there is conservation of some functional pathways, we found that key age-related pathways either lost rhythmicity or showed significant changes in their timing or phase of expression. Our observations identify age-associated changes in circadian gene expression as well as temporal shifts in gene expression. We propose that the age-associated changes in circadian clock output leads to reduced predictive homeostasis and an increased reliance on reactive pathways in response to stressors. This loss of predictive homeostasis pathways is part of a new concept in aging that will contribute to decreased resilience and frailty (Figure 7I). We suggest that circadian clock output be considered as a new hallmark of aging and that the circadian clock and its transcriptional program may serve as a new therapeutic target for improving tissue and systemic health across the lifespan.

Age-related changes in gene expression have been providing significant insights into the understanding of the etiology of aging (Aging Atlas Consortium, 2021; Schaum et al., 2020; Shavlakadze et al., 2019; Tacutu et al., 2018; Zahn et al., 2007). Interestingly, clock genes were among the most differentially expressed across tissues with age (Schaum et al., 2020). We leveraged our time course data to ask whether there are clusters of aging genes and pathways that are unique to specific phases of the day. The discovery that there are tissue-specific and time-domain aging clusters is novel and includes many aging hallmark pathways. One example is that genes related to autophagy were upregulated with age in the rest phase in the hypothalamus but in the active phase in skeletal muscle. The increase in autophagy-related DEGs occurred as the circadian regulation of autophagy-related genes was lost, consistent with emerging evidence highlighting the importance of the circadian control of autophagy in healthy aging (Ulgherait et al., 2021). By considering the time of day, the DEGs provide a new, more precise insight into the complexity of aging.

The age-dependent circadian decline can result from internal and external factors. A better understanding of the role of the circadian clock in aging physiology can offer new strategies for interventions. As environmental factors such as light, food intake, and physical activity are known to influence the circadian system, lifestyles that conform to circadian rhythms may represent an attractive non-pharmacologic regimen to slow down aging and improve healthspan. For example, time-restricted feeding regimen extends healthspan at least in part by improving the circadian transcriptome (Chaix et al., 2019; Lundell et al., 2020). It should be noted that our data demonstrated a rapid decline from Aged to Old. Thus, the middle age period may represent a window of opportunity for early intervention that considers circadian concepts in healthy lifestyle choices. Mechanistically, enhanced circadian function improves cell and systemic homeostasis which in turn can delay and mitigate damage accumulation, frailty phenotypes, increase resilience, and extend healthspan.

In summary, this aging circadian Resource provides a rich dataset highlighting the impact of both age and time of day on the transcriptome in multiple tissues. We suggest that the age-related changes in clock output are linked to a loss of resilience. This dataset will help foster collaborations across multiple disciplines. We recognize the potential for sex differences in the circadian transcriptome, especially given known sex differences in aging (Austad and Fischer, 2016; Lemaître et al., 2020) warranting additional work in female mice. Future work to determine if the age-related changes in circadian clock output and function can be reversed by known pro-longevity interventions should be explored.

## Acknowledgements

We thank the members of the Esser, Liu, Bryant, Gumz, and Huo laboratories for their work with the circadian collection, sample preparations, and lively discussions while developing this dataset and manuscript. We also thank Drs. Yanping Zhang and David Moraga from the University of Florida Interdisciplinary Center for Biotechnology Research (ICBR) for library preparation and sequencing (RRID:SCR_019152) and the UF High performance computing cluster. This work was supported by grants (R01HL141343 to BPD; R01HL142776 and R01HL142887 to AJB; R56DK128271 and R01DK109570 to MLG; R01NS054794 to ACL; R01HL153042 to BPD and KAE, and R01AR079220 to KAE) and support from the UF NIH Claude D. Pepper Older Americans Independence Center (P30AG028740) for ACL and KAE and the Office of Research Affairs, UF College of Medicine.

## Author Contributions

Conceptualization, CAW, MAG, ACL, MLG, AJB, and KAE; Methodology, CAW, LM, ZH; Investigation, CAW, XZ, MAG, CMD, ARM, MME, EE, EAS, BPD; Writing – Original Draft, CAW, MAG, ACL, KAE; Writing – Review & Editing, All authors reviewed and approved the final version of this manuscript; Funding Acquisition, BPD, AJB, MLG, ZH, ACL, and KAE; Resources, ACL, MLG, AJB, ZH, and KAE; Supervision, ACL, MLG, ZH, AJB, and KAE.

## STAR METHODS

### EXPERIMENTAL MODEL AND SUBJECT DETAILS

#### Animal Care and Use

6 month, 18 month, and 27 month old C57BL/6Nia male mice (n=24/age) were used to assess circadian robustness and molecular clock transcriptional output. All procedures were approved by the University of Florida Institutional Animal Care and use Committee, in accordance with AAALAC guidelines. Mice were ordered from the National Institute on Aging (NIA) colony 2 months prior to collections. Following a two week quarantine, animals were allowed a two week acclimation period to ensure entrainment to the 12h:12h light/dark cycle (lights on: 6am). Prior to tissue collection, animals were released into constant darkness in light-tight circadian cabinets (Actimetrics, Wilmette, IL, USA). To assess the circadian transcriptome in each age, collections occurred every 4h for 48h, starting at CT18. Mice were euthanized by cervical dislocation under dim red light. 2 animals for each age were collected at each time point.

#### RNA Isolation and Library Preparation

RNA was isolated from the hypothalamus, kidney, lung, gastrocnemius, adrenal glands, and heart using a modified TRIzol extraction procedure as previously described (Hodge et al., 2019; Terry et al., 2018). Briefly, frozen tissues were lightly ground in a mortar and pestle constantly submerged in liquid nitrogen. Frozen tissue between 10 and 100 mg was placed into RNase-free tubes containing TRIzol and RNase-free stainless-steel beads ranging in diameter from 0.9 to 2.0mm (NextAdvance, Troy, NY, USA). Tissues were homogenized using a bullet blender at 4°C. Chloroform was used to separate the phases, and the RNA-containing aqueous phase was subjected to a modified column purification kit using RNeasy Mini RNA extraction kit for all tissues except the hypothalamus, prepared with RNesay Micro RNA extraction kit (Qiagen, Germantown, MD, USA). RNA was DNase treated, and purified RNA was checked for integrity using an Agilent Bioanalyzer, all samples had an RNA integrity number above 8.0. Poly-A selected RNAseq libraries were prepared using Illumina mRNA Prep kit (Illumina, San Diego, CA, USA). Libraries were pooled to equal molarity and sequenced on an Illumina NovaSeq (2×100bp) to achieve a minimum of 40M reads per sample. FastQ files were downloaded to the University of Florida HiPerGator computing cluster. Raw FastQ files, counts, and normalized counts per million data were uploaded to GEO: GSE201207.

### QUANTIFICATION AND STATISTICAL ANALYSIS

#### Bioinformatics and data preprocessing for RNA-Sequencing data

Reads from the RNA-Sequencing data were aligned to the *Mus musculus* genome assembly GRCm38 (mm10) using the HISAT2 software. Duplicated aligned reads were marked and removed using the Picardtools software. The gene expression count data were extracted using the HTseq software. The raw count data were normalized to cpm (counts per million reads), followed by log2 transformation (i.e., log2(cpm+1) where 1 is the pseudo count to avoid log2(0)). For each tissue, we filtered out genes with mean log2(cpm+1) < 1. These nonexpressed genes were likely to be false positives, and thus removing them could help reduce the multiple testing burden in the later statistical analysis. All data preprocessing was performed using R software.

#### Circadian rhythmicity analysis

To identify genes showing circadian rhythmicity patterns for each age group within each tissue type, we deployed the cosinor model implemented in the diffCircadian software. To be specific, the cosinor model assumes the following relationship between the expression level and

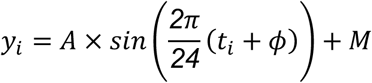

Where *i* (1 ≤ *i* ≤ *n*) is the sample index; *n* is number of samples in each age group within each tissue type; *y_i_* is the log2 transformed cpm value for sample *i*; *t_i_* is the circadian time for sample *i*; *A, ϕ*, and *M* denote the amplitude, phase, and MESOR (Midline Estimating Statistic of Rhythm) of the sinusoidal curve; the period was fixed at 24-hour. The goodness of fit of the cosinor model was accessed via the coefficient of determination (*R^2^*), and the p-value was determined by the finite-sample-corrected likelihood ratio test [cite]. Raw p-value<0.01 was used as the statistical significance cutoff to declare rhythmically expressed genes (REGs). To further compare the phase concordance between age groups, the Watson’s Two-Sample Test of Homogeneity was adopted. This comparison was performed for each pair of age groups, and for each tissue, respectively.

#### Differential expression analysis associated with aging

We first created 4 time domain of day groups: including (i) Active Phase: CT18,22,42,46; (ii) Activity Offset: CT22,26,46,50; (iii) Rest Phase: CT30,34,54,58; and (iv) Activity Onset: CT34,38,58,62. To detect genes showing increasing/decreasing expression values with respect to the ordinal age groups (Young to Aged to Old), we employed negative binomial models implemented in R software edgeR package, which is specifically designed for RNA-Sequencing count data. In this model, the expression value of a gene was the outcome variable, the ordinal age groups were coded as the continuous predictor (0: Young group; 1: Aged group; 2: Old group). Multiple testing was corrected by the Benjamini-Hochberg method, in which the raw p-values were converted to q-values (FDR-adjusted p-values), where FDR stands for the false discovery rate. This analysis was performed for each time of day group within each tissue, respectively. Q-value<0.05 was used as the statistical significance cutoff to declare differentially expressed genes (DEGs).

#### Variability analysis associated with aging

To identify genes showing increasing/decreasing variability in expression values with respect to the ordinal age groups (Young to Aged to Old), we used *DiffVar* function in the *missMethyl* package (Phipson and Oshlack, 2014). The variability for each sample of a gene in one age group was calculated using absolute deviations. A linear model with empirical Bayes shrinkage was then fitted, where the variability was the outcome variable, the ordinal age groups were coded as the continuous predictor (0: Young group; 1: Aged group; 2: Old group). P-value<0.01 was used as the statistical significance to declare variably expressed genes.

#### Pathway analysis

To examine the function annotation of the putative REGs (raw p-value < 0.01), DEGs (q-value < 0.05), and variably expressed genes (raw p-value < 0.01), pathway enrichment analyses were performed using Ingenuity Pathway Analysis software (REGs), and both g:Profiler (Raudvere et al., 2019) and DAVIDV6.9 (Huang et al., 2009) for the DEGs.

To further compare and integrate the REGs’ pathway information (i.e., p-values) across all three age groups, an adaptively weighted Fisher’s method was adopted, which is a metaanalysis to examine whether a pathway is enriched in at least one age group. Unlike the regular Fisher’s method that assigns equal weight for all three age groups, the adaptively weighted method searches for the optimal binary adaptive weight (1 for enriched and 0 for not enriched) for each age group (Young/Aged/Old) given a pathway. These binary adaptive weights can capture the similarities across the three age groups. For instance, an adaptive weight of (1,1,1) indicates a pathway is enriched in all three age groups; and an adaptive weight of (1,0,0) indicates a pathway is enriched only in the young group. A meta-analyzed p-value was also reported, followed by multiple comparison correction using the Benjamini-Hochberg method. This analysis was performed for each tissue type, respectively.

## Supplemental Information Title and legends

Supplemental Table 1: Circadian F-test output data. One tab for each tissue, all genes passing filter criteria are included. Amp – circadian amplitude; phase time of gene peak plus 6h; peakTime circadian time at which the gene has highest expression; basal mean expression level across all 12 time points; R^2^ – R^2^ goodness of fit value for cosine curve; p value significance of goodness of. fit; qvalue multiple comparison adjusted p value for significance of fit. Rhythmically expressed genes were those with a raw p-value <0.01.

Supplemental Table 2: Age-specific and shared REGs across ages in the hypothalamus. Genes with a raw p-value <0.01 were considered rhythmic for a given age and genes with a raw p-value >0.10 were considered non-rhythmic (red shading).

Supplemental Table 3: Age-specific and shared REGs across ages in the lung. Genes with a raw p-value <0.01 were considered rhythmic for a given age and genes with a raw p-value >0.10 were considered non-rhythmic (red shading).

Supplemental Table 4: Age-specific and shared REGs across ages in the kidney. Genes with a raw p-value <0.01 were considered rhythmic for a given age and genes with a raw p-value >0.10 were considered non-rhythmic (red shading).

Supplemental Table 5: Age-specific and shared REGs across ages in the skeletal muscle. Genes with a raw p-value <0.01 were considered rhythmic for a given age and genes with a raw p-value >0.10 were considered non-rhythmic (red shading).

Supplemental Table 6: Age-specific and shared REGs across ages in the heart. Genes with a raw p-value <0.01 were considered rhythmic for a given age and genes with a raw p-value >0.10 were considered non-rhythmic (red shading).

Supplemental Table 7: AW-Fisher analysis of significantly enriched pathways across ages from the rhythmically expressed genes in the hypothalamus. A value of 1 denotes significant enrichment for an age and 0 denotes no significant enrichment. Full pathway data from each age is also included as individual tabs.

Supplemental Table 8: Gene lists and accompanying gene ontology data for the REGs within a 4h window of the day from the hypothalamus. One tab for each 4h window from each age. Correspond with Panels C-E from Figure 2.

Supplemental Table 9: DEGs from the hypothalamus at each time domain with ordinal slope, p-value, and q-value. Accompanying biological processes are also included on each tab for each gene list.

Supplemental Table 10: AW-Fisher analysis of significantly enriched pathways across ages from the rhythmically expressed genes in the lung. A value of 1 denotes significant enrichment for an age and 0 denotes no significant enrichment. Full pathway data from each age is also included as individual tabs.

Supplemental Table 11: Gene lists and accompanying gene ontology data for the REGs within a 4h window of the day from the lung. One tab for each 4h window from each age. Correspond with Panels C-E from Figure 3.

Supplemental Table 12: DEGs from the lung at each time domain with ordinal slope, p-value, and q-value. Accompanying biological processes are also included on each tab for each gene list.

Supplemental Table 13: AW-Fisher analysis of significantly enriched pathways across ages from the rhythmically expressed genes in the kidney. A value of 1 denotes significant enrichment for an age and 0 denotes no significant enrichment. Full pathway data from each age is also included as individual tabs.

Supplemental Table 14: Gene lists and accompanying gene ontology data for the REGs within a 4h window of the day from the kidney. One tab for each 4h window from each age. Correspond with Panels C-E from Figure 4.

Supplemental Table 15: DEGs from the kidney at each time domain with ordinal slope, p-value, and q-value. Accompanying biological processes are also included on each tab for each gene list.

Supplemental Table 16: AW-Fisher analysis of significantly enriched pathways across ages from the rhythmically expressed genes in the skeletal muscle. A value of 1 denotes significant enrichment for an age and 0 denotes no significant enrichment. Full pathway data from each age is also included as individual tabs.

Supplemental Table 17: Gene lists and accompanying gene ontology data for the REGs within a 4h window of the day from the skeletal muscle. One tab for each 4h window from each age. Correspond with Panels C-E from Figure 5.

Supplemental Table 18: DEGs from the skeletal at each time domain with ordinal slope, p-value, and q-value. Accompanying biological processes are also included on each tab for each gene list.

Supplemental Table 19: AW-Fisher analysis of significantly enriched pathways across ages from the rhythmically expressed genes in the heart. A value of 1 denotes significant enrichment for an age and 0 denotes no significant enrichment. Full pathway data from each age is also included as individual tabs.

Supplemental Table 20: Gene lists and accompanying gene ontology data for the REGs within a 4h window of the day from the heart. One tab for each 4h window from each age. Correspond with Panels C-E from Figure 6.

Supplemental Table 21: DEGs from the heart at each time domain with ordinal slope, p-value, and q-value. Accompanying biological processes are also included on each tab for each gene list.

Supplemental Table 22: Lists of Variably Expressed Genes in each tissue with age. Change in variability and accompanying raw p or multiple comparison adjusted q values. Accompanying biological process output for each gene list is presented as well.

## FIGURES

Supplemental figure 1: Age-related changes in circadian transcriptome and non-circadian gene expression in the adrenal gland.

Supplemental figure 2: Age-specific REGs. Heatmaps demonstrating REGs unique to each age within a tissue.

Supplemental figure 3: Core clock gene expression. Circadian F-test p values were negative log transformed and presented for each core clock gene at each age and for each tissue. A horizontal line is present for each gene at Y=2, our chosen cut off for significant circadian rhythmicity.

