## Supplementary figures and images for "Defining the age-dependent and tissue-specific circadian transcriptome in male mice"

### Figure S1

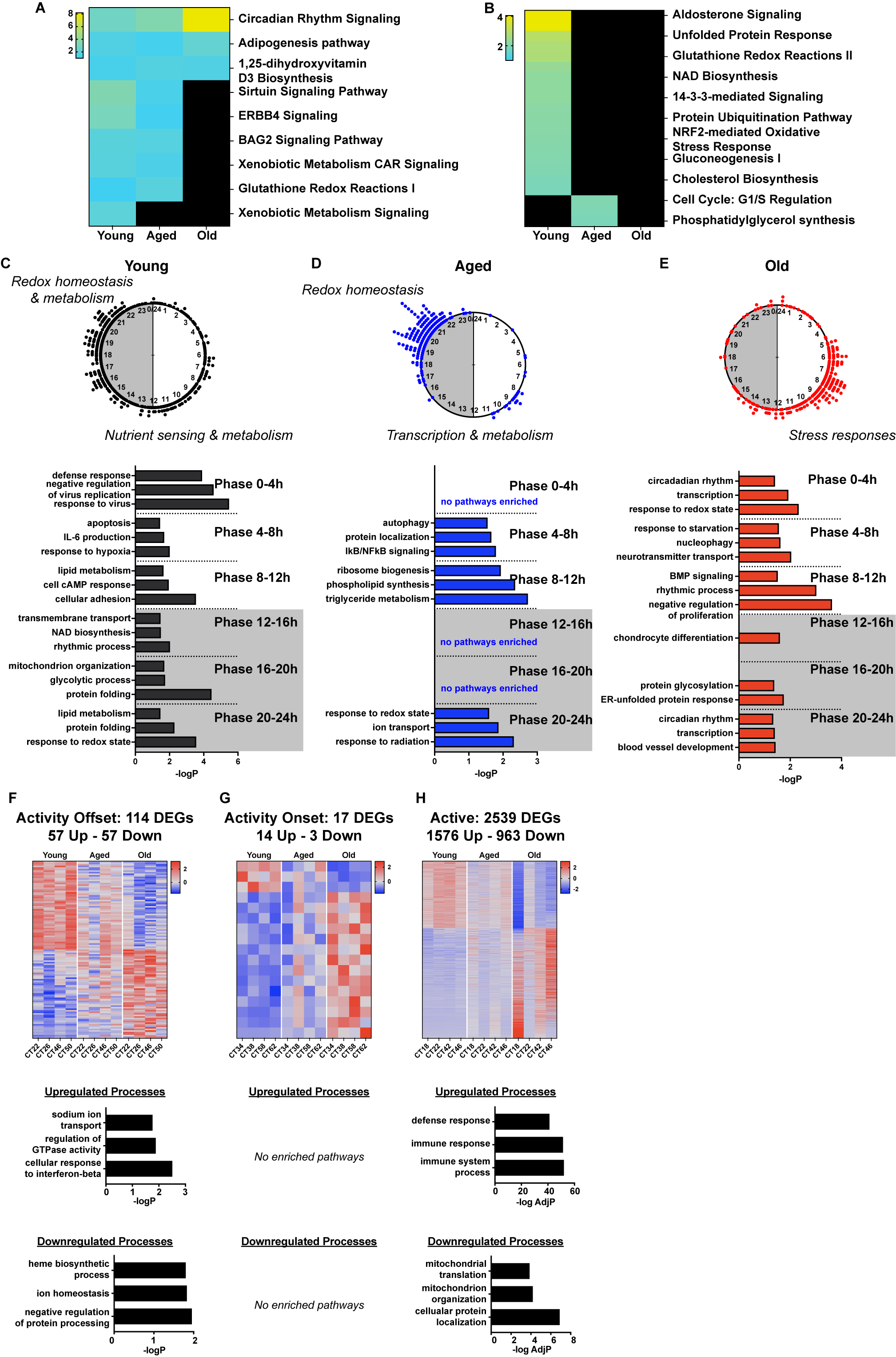

### Figure S2

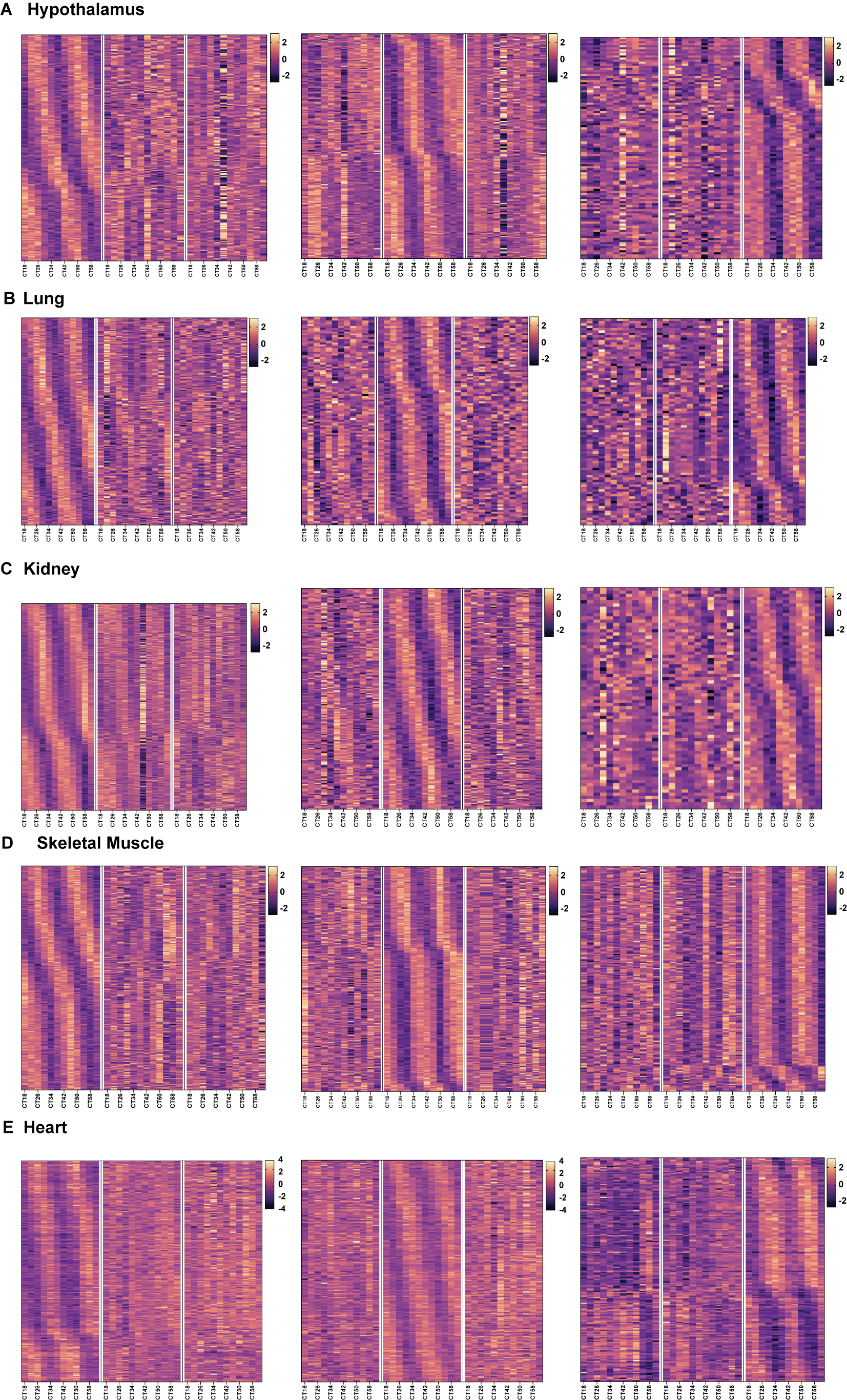

### Figure S3

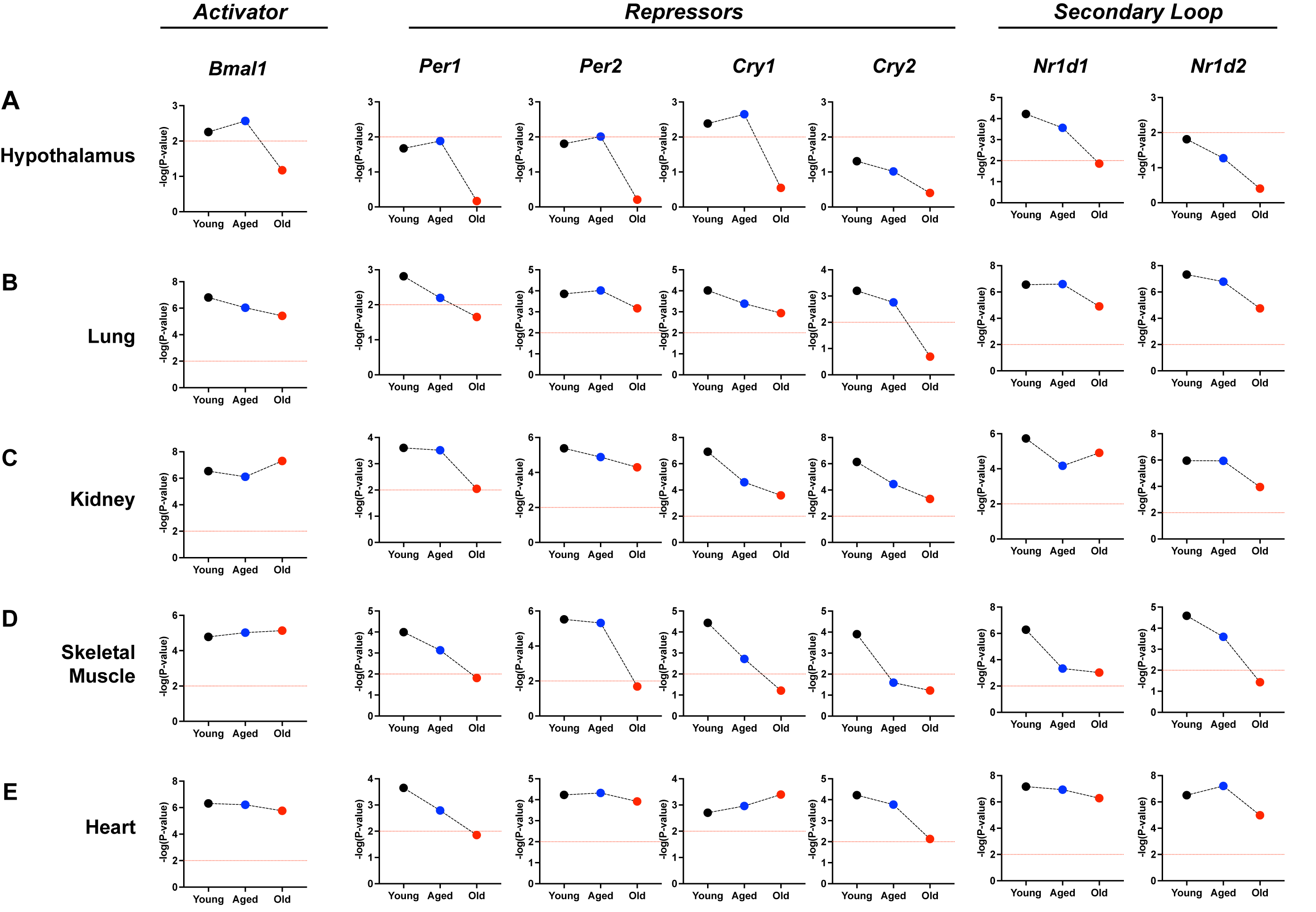
